# *Bacillus sp.* RZ2MS9, a tropical PGPR, colonizes maize endophytically and alters the plant’s production of volatile organic compounds both independently and when co-inoculated with *Azospirillum brasilense* Ab-V5

**DOI:** 10.1101/2021.01.04.425352

**Authors:** Jaqueline Raquel de Almeida, Maria Letícia Bonatelli, Bruna Durante Batista, Natalia Sousa Teixeira-Silva, Mateus Mondin, Rafaela Cristina dos Santos, José Maurício Simões Bento, Carolina Alessandra de Almeida Hayashibara, João Lúcio Azevedo, Maria Carolina Quecine

## Abstract

*Bacillus* spp. are among the most efficient known plant growth-promoting rhizobacteria (PGPR). The PGPR *Bacillus* sp. strain RZ2MS9 is a multi-trait maize growth promoter previously isolated from guarana plants cultivated in the Amazon rainforest. However, there are several aspects of its interaction with the host that need further investigation. To achieve effective performance of microbial inoculants in crop production, it is necessary to monitor the plant’s colonization by a PGPR and to assess the potential synergy among beneficial strains. Here, we obtained a stable mutant of RZ2MS9 labelled with *green fluorescent protein* (RZ2MS9-GFP). We verified that the insertion of the plasmid did not affect either bacterial growth nor its ability to promote maize growth *in vitro*. Using fluorescent microscopy and qPCR, we demonstrated that RZ2MS9-GFP successfully colonizes maize’s roots and leaves endophytically. Subsequently, we evaluated whether RZ2MS9 has a synergistic effect on plant growth promotion when co-inoculated with *Azospirillum brasilense* Ab-V5, a commercial inoculant for maize. The two strains combined enhanced maize’s roots and shoots dry weight by 50.8% and 79.6%, respectively, when compared to the non-inoculated control. In addition, we used co-inoculation experiments in glass chambers to analyze the plant’s Volatile Organic Compounds (VOCs) production during the maize-RZ2MS9 and maize-RZ2MS9-Ab-V5 interaction. We found that the single and co-inoculation altered maize’s VOCs emission profile, with an increase in the production of indoles in the co-inoculation. Collectively, these results increase our knowledge about the interaction between the tropical PGPR *Bacillus* sp. RZ2MS9 and maize, and provide a new possibility of combined application with the commercial inoculant *A. brasilense* Ab-V5.

**Importance:** *Bacillus* sp. RZ2MS9 is a PGPR, previously isolated from guarana plants cultivated in the Brazilian Amazon, which successfully promotes the growth of maize and soybean plants. To improve our knowledge about the interaction between this very promising PGPR and maize, we labelled RZ2MS9 with *gfp* and monitored it’s maize colonization. The transformation did not affect either RZ2MS9 growth nor its ability to promote maize growth *in vitro*. We demonstrated that RZ2MS9 colonizes endophytically maize’s roots and leaves. We also verified that the co-inoculation of RZ2MS9 and *Azospirillum brasilense* Ab-V5, a known commercial maize inoculant enhanced maize’s roots and shoots growth. Moreover, the co-inoculation altered the maize’s volatile organic compounds, increasing the production of indoles, that is related with decreased upon the reduction of fertilization. Certainly, our research contributed with better *Bacillus* sp. RZ2MS9 – maize interaction understanding and also provided new information concerning RZ2MS9 activity when applied with *A. brasilense* Ab-V5.

## INTRODUCTION

Plant growth-promoting rhizobacteria (PGPR) are beneficial bacteria which are able to establish a symbiotic or nonsymbiotic association with plants in the rhizosphere (Miransari, 2016). A PGPR can colonize the plant’s interior tissues and thrive as endophytes. The understanding of this process in plants of agricultural importance can be used to increase crop production (Compant et al., 2008).

Once associated with the host, PGPR can improve the plant growth through many mechanisms, such as producing phytohormones (Patten and Glick, 1996), fixing biological nitrogen (Hurek et al., 2002), solubilizing phosphorus (Rodriguez et al., 2004; Vikram and Hamzehzarghani, 2008), improving nutrient absorption (Kraiser et al., 2011), producing antimicrobial metabolites (Raaijmakers et al., 2002), triggering the induction of plant systemic resistance (Bakker et al., 2007), as well as modulating plant growth through the production of volatile organic compounds (VOCs) (Fincheira et al., 2018). It is known that PGPR can alter the plant’s VOC signaling (Santoro et al., 2015; Cappellari et al., 2017). However, most studies focus on how the emission of microbial VOCs interfere with plant development (Chowdhury et al., 2018), and not on how the plant’s VOC profile changes upon microbial inoculation.

Among the PGPR, members of the Gram-positive endospore-forming *Bacillus* group are the most commonly reported. Because of the advantages of using *Bacillus* as inoculants, such as high cell viability and prolonged shelf-life when in formulation, this group is frequently commercialized (Akinriniola et al., 2018). The *Bacillus* sp. strain RZ2MS9 is a rhizobacterium isolated from guarana plants (*Paullinia cupana*) cultivated in the Brazilian Amazon (Batista et al., 2016). The strain appears to have no specificity for the host plants, successfully promoting the growth of important crops, as maize and soybean (Batista et al., 2018). The inoculation of RZ2MS9 increased substantially the dry weight of maize shoots and roots compared with the non-inoculated controls. In soybean, RZ2MS9 also increased the shoot and root’s dry weight (Batista et al., 2018). In addition, plant growth-promoting mechanisms such as indole acetic acid production (IAA), biological fixation of nitrogen, and phosphate solubilization have been detected *in vitro* in *Bacillus* sp. RZ2MS9 (Batista et al., 2018). Several genes involved in these traits were identified in the strain’s genome (Batista et al., 2016; Bonatelli et al., 2020). It is commonly reported that a single PGPR can exhibit more than one of the above-mentioned plant growth-promoting mechanisms (Ahmad et al., 2008). However, there is an increasing trend to use products based on microbial consortium, with the aim to exploit their complementary or even synergistic interactions (Bradáčová et al., 2019). For instance, the co-inoculation of *Azospirillum* spp. with other PGPR was superior in increasing the rice growth and yield in contrast with single inoculation (Amutha et al., 2009). The co-inoculation of two PGPR, *Paenibacillus polymyxa* and *Bacillus megaterium*, and of three rhizobia, IITA-PAU 987, IITA-PAU 983 and CIAT 899, in different combinations, showed a synergistic effect on the growth of common bean (*Phaseolus vulgaris* L.) (Korir et al., 2017).

Brazil is the world’s third largest maize producer, with an expected production of 95 million tons in the Market Year 2019/2020 (USDA, 2019). The commercial use of *Azospirillum brasilense* strains Ab-V5 and Ab-V6 on maize crops in Brazil has grown exponentially since 2010 (Fukami et al., 2017). Inoculants formulated with *A. brasilense* can reduce nitrogen application by up to 25% with increases in maize yield of up to 30%. The bacterial traits that best explain its beneficial association with cereals are nitrogen fixation, phytohormones production, mitigation of abiotic stresses, and control of plant pathogens (Pereira et al., 2020).

Considering the beneficial effects of both *Bacillus* sp. RZ2MS9 and *A. brasilense* on maize development, we hypothesized that their combined application would provide a more robust effect on maize growth as compared to the single RZ2MS9 application. Thus, this work aimed to monitor the colonization of *Bacillus* sp. RZ2MS9 in maize plants and to test the effect of co-inoculating *Bacillus* sp. RZ2MS9 and *A. brasilense* Ab-V5 on maize’s growth and production of VOCs.

## MATERIAL AND METHODS

### Bacterial strains and growth conditions

*Bacillus* sp. RZ2MS9, a plant growth-promoting rhizobacteria previously isolated from guarana (*Paullinia cupana*) (Batista et al., 2016), and its transformant, the GFP-tagged strain, were routinely grown in Luria-Bertani (LB) medium (Sambrook et al., 1998) at 28 °C, using appropriated antibiotic when necessary. *A. brasilense* Ab-V5, a commercial maize inoculant (Hungria et al., 2010), was routinely grown in DYGS medium (Rodriguez et al., 2004) at 28°C. All strains are stored in 20% glycerol at −80°C. The integrative plasmid pNKGFP (Ferreira et al., 2008) was propagated and isolated from *E. coli* DH5α-pir and purified with QIAprep spin miniprep kit (Qiagen) according to the manufacturer’s recommendations.

### Development of stable GFP-tagged RZ2MS9

#### Transformation of *Bacillus* sp. RZ2MS9 by electroporation

RZ2MS9 transformants were obtained by electroporation according to the protocol described by Schurter et al. (1989), with modifications. Briefly, one single colony of RZ2MS9 was inoculated into 10 ml of LB amended with glycine (0.1%) and incubated overnight in an incubator shaker at 150 rpm and 28°C. The culture was 100-fold diluted (optical density - OD_550nm_ = 0.01) in LB with glycine (0.1 %) and incubated until it reached the OD_550nm_ = 0.2. The bacterial cells were harvested by centrifugation and resuspended twice in 1/40 of the volume in ice-cold electroporation buffer (400 mM sucrose, 1 mM MgCI_2_, 7 mM phosphate buffer, pH 6.0). Then, the bacterial cells were resuspended into 2.5 ml of the electroporation buffer and 800 μl of the suspension were distributed into precooled 2 mm cuvettes. The pNKGFP plasmid (30 ng) was also added into the cuvettes and kept for 10 min at 4°C. The electroporation was performed using 25 μF, and 200 Ω. After electroporation, the cuvettes were maintained for 10 min at 4°C, diluted into 1.2 ml LB and incubated for 2 h at 28°C under agitation (150 rpm). After this period, the bacterial suspension was spread onto LB plates supplemented with kanamycin (50 μg.ml^−1^) and incubated at 28°C for 24 h. One colony was randomly selected, named RZ2MS9-GFP, grown in LB broth medium containing kanamycin (50 μg.ml^−1^), and preserved in 20% glycerol at −80°C for further studies.

#### Molecular confirmation of the transformation

One single colony of RZ2MS9 wild-type (wt) and one of the transformant RZ2MS9-GFP were inoculated into 5 ml of LB medium and maintained overnight at 28°C under agitation (150 rpm). RZ2MS9-GFP was also grown in LB medium supplemented with kanamycin (50 μg.ml^−1^) under the same conditions. Bacterial cells were harvested by centrifugation and the genomic DNA was extracted using the DNeasy^®^ blood and tissue kit (Qiagen) following the manufacturer’s recommendations. The DNA integrity was verified in agarose gel (1%) stained with 0.5x SYBR^®^ Green (Invitrogen^®^) and quantified in NanoDrop™ spectrophotometer (NanoDrop Technologies).

To confirm the RZ2MS9-GFP transformation, we amplified the internal region of the plasmid using the primers: PPNKF (5 ‘CCTTCATTACAGAAACGGC 3’) and PPNKRII (5 ‘GGTGATGCGTGATCTGATCC 3’) (Quecine et al., 2012). The pNKGFP plasmid was used as a positive control, while the RZ2MS9 wt DNA and a DNA-free water were used as negative controls. The reaction was performed with 0.75 μl of MgCl_2_ (25 mM), 0.5μl of dNTP (10 μM), 2.5 μl of 10X Buffer, 0.5 μl of each primer (10 μM), 0.3 μl of Taq DNA polymerase, 19 μl of water, yielding a final volume of 25μl per reaction. The PCR program consisted of an initial denaturation step at 94°C for 4 min, followed by 35 cycles of denaturation at 94°C for 30s, annealing at 58°C for 45 s, and extension at 72°C for 30 s. The final step was a 10 min extension at 72°C. The amplified products were separated by electrophoresis in an agarose gel (1%) stained with 0.5x SYBR Green (Invitrogen).

#### Influence of the transformation on the bacterial growth

To evaluate whether the plasmid pNKGFP can affect the transformant growth, the RZ2MS9 wt and the RZ2MS9-GFP were simultaneously grown in 50 ml of LB medium (the LB medium was supplemented with kanamycin 50 μg.ml^−1^ for the transformant growth). The OD_550nm_ of both bacterial cultures was standardized to 0.2 using LB medium for a final volume of 50 ml. The flasks were then incubated at 28°C under agitation (150 rpm) and the OD_550nm_ was measured every 2, for 10 h, and then at 24 h and at 48 h after inoculation (h.a.i). The experiment was performed using four replicates.

#### Influence of the transformation on the bacterium-maize interaction

We evaluated the effect of the *Bacillus* sp. RZ2MS9 wt and of the RZ2MS9-GFP on the initial development of maize seedlings. The maize seeds, cultivar Dupont Pioneer^®^ P4285H, were washed twice in distilled water and then immersed in the respective bacterial solutions (OD_600_: 0.12) for 30 min. The control treatment consisted of immersing the washed seeds in LB medium without any bacterial growth.

The treated seeds and the control were seeded onto wet germination paper towels, placed into Petri dishes, and incubated in the dark at 28°C for 7 d. The assay was performed using 10 seeds per plate and four replicates (plates) per treatment.

The root system growth was assessed using a scanner. The images of 12 seedlings per treatment were captured at 400 dpi (dots per inch) resolution with an Epson^®^ Expression 11000XL scanner and analyzed using the software WinRHIZO Arabidopsis (Regent Instruments Inc., Quebec, Canada). The parameters of root system development measured were root volume (cm^3^), axial root (cm), lateral root (cm), surface area (cm^2^), diameter (mm) and length (mm).

### Maize colonization by RZ2MS9-GFP

Maize colonization by RZ2MS9-GFP was monitored by applying the transformant culture to the seeds and growing the plants under greenhouse conditions. Seed inoculation was performed according to Batista et al. (2018), with modifications. Briefly, culture cells were obtained by growing the RZ2MS9-GFP in 100 ml of LB supplemented with kanamycin (50 μg.ml^−1^) at 28°C in a shaker incubator (150 rpm) until it reached the late log phase. The cells were harvested by centrifugation at 4,500 g for 15 min, washed with phosphate buffer saline - PBS (pH 6.5), and adjusted to a cell density of 10^8^ colony-forming units (CFU).ml^−1^. The seeds, cultivar Dupont Pioneer^®^ P4285H, were immersed into the bacterial solution for 30 min and then sown. The control treatment consisted of immersing the seeds into PBS for 30 min. The treated seeds and the control were seeded in pots containing 1.6 kg of the thick, branny substrate Bioplant^®^ (http://www.bioplant.com.br/), which is composed of peat, correctives, vermiculite, charcoal and pine bark (Bioplant Agrícola Ltda.). The pots were kept at 28°C and daily irrigated in a greenhouse located at the “Luiz de Queiroz” College of Agriculture, University of São Paulo, Piracicaba – SP, Brazil (22° 42′ 30″ S and 47° 38′ 30″ W). Six plants per treatment were collected at 15 and at 30 d after germination (DAG) for further analyzes.

### Quantitative PCR (qPCR)

For quantification by qPCR, the plants collected (both at 15 and at 30 DAG) were immediately stored at −80°C for later DNA extraction. The total DNA was extracted using the DNeasy^®^ Plant Mini Kit (Qiagen) following the manufacturer’s instructions. The DNA integrity was verified using agarose gel (1%) electrophoresis stained with 0.5x SYBR^®^ Green (Invitrogen^®^) and observed under UV light. The DNA quantification was performed using the NanoDrop^™^ spectrophotometer (NanoDrop Technologies).

For RZ2MS9-GFP quantification, we used the primers PPNKF and PPNKRII (Quecine et al., 2012). The qPCR analysis was performed in 12 μl final volume, containing 6.25 μl of the Platinum^®^ qPCR superMix-UDG (Invitrogen), 0.25 μl of each primer - PPNKF and PPNKRII (10 μM) - and 0.25 μl of Bovine Serum Albumin (BSA). Aliquots of the master mix (7 μl) were distributed in the wells of a 96-well plate, and 5 ng of DNA was added as a PCR template. The qPCR cycles consisted of a denaturation step at 95°C for 2 min, 40 cycles at 95°C for 30 s, and a final step at 62°C for 15 s. The quantification was performed in the iCycler iQ real-time PCR instrument (BioRad Laboratories Inc.). Three biological replicates and two technical replicates were used.

The standard curve was obtained for each run using a known copy number (10^4^ to 10^10^) of the linearized plasmid pNKGFP. The bacterial density (CFU per nanogram of total plant DNA) was estimated by interpolation with the standard curve (Quecine et al., 2012).

### Fluorescence Microscopy (FM)

Immediately after the plant sampling, both at 15 and 30 DAG, root systems and shoots were cut and observed using the epifluorescence microscope Axiophot II (Zeiss, Germany). The filter sets (excitation/emission) used were 365/397 nm for blue, 450/515 nm for green, and 546/590 nm for red. The images were digitalized through a PCO CCD camera using the ISIS Metasystems software (Metasystems, Germany).

### Assessment of maize growth promotion with co-inoculation of RZ2MS9 and Ab-V5

To evaluate whether the Ab-V5 improves the beneficial effect of the RZ2MS9 on maize growth, cultures of both bacteria were obtained in a final concentration of 10^8^ CFU.ml^−1^ as previously described (the DYGS medium was used for the strain Ab-V5). The maize seeds, cultivar Dupont Pioneer^®^ P4285H, were immersed into the bacterial solutions for 30 min and then sown. The control treatment consisted of immersing the seeds into PBS for 30 min.

The treated seeds and the control were seeded in pots containing 1.6 kg of the thick, branny substrate Bioplant^®^ (http://www.bioplant.com.br/) as previously described. Four seeds were sown per pot and thinning was performed at 8 DAG, leaving two plants per pot. At 15 DAG, one plant per pot was collected for the first assessment. The second assessment was performed at 30 DAG with the remaining plants. In total, six plants were collected per treatment in each assessment. The roots were separated from the shoots, washed and kept in pots containing alcohol (70%). The height of the plant shoots and diameters were measured, and then the shoots were relocated in paper bags. The shoots’ dry weight was measured in an analytical balance after oven-drying at 70°C for 5 d.

The root systems were scattered in a clear layer of water in a tray (30 cm by 20 cm), and the images were captured at 400 dpi with an Epson^®^ Expression 11000XL professional scanner system. The obtained images were analyzed using the software WinRHIZO Arabidopsis (Regent Instruments Inc., Quebec, Canada), as previously described. In addition, the ten diameter classes provided by the software were simplified in only two: axial and lateral root length (Trachsel et al., 2009). Therefore, we considered the root portions with a diameter of less than or equal to 0.5 mm for lateral root length measurement, and the root portions with a diameter greater than 0.5 mm for axial roots length measurement. After, the roots were relocated in paper bags and their dry weight was measured as previously described.

### Volatile-collection system and analyzes

The volatile organic compounds (VOCs) emitted by 15 days-old maize plants were collected in a system described by Turlings et al. (1998). The maize VOCs were collected from plants inoculated with RZ2MS9, from plants co-inoculated with RZ2MS9 and Ab-V5, and from non-inoculated plants (control). Six plants per treatment were enclosed in glass chambers and connected to the ARS Volatile Collection System (Analytical Research Systems, Gainesville, FL, USA) through PTFE (polytetrafluoroethylene) hoses. Clean humid air was first pushed into each glass chamber with flow of 0.3 L.min^−1^. A column with 30 mg of Hayesep-Q^®^ adsorbent polymer (Alltech Associates, Deerfield, IL, USA) was connected to each glass chamber. The chambers were linked with hoses to a vacuum pump that was pulling the air. The VOCs collection assay was kept for 12h (from 7:00 am to 7:00 pm) in a room with controlled temperature (25 ± 1°C), relative humidity (60 ± 10%), and photoperiod (12 h of light/12 h of dark). After, we eluted the polymer columns with 150μl hexane and added 10 μl of nonyl acetate (10 ng.μl^−1^) into each sample as an internal standard. Samples were stored in a freezer at −30°C until further analysis.

Two microliters of the samples were injected into a GC-FID, Shimadzu 2010 chromatrograph to quantify the maize’s emitted VOCs, while 1 μl aliquot was injected into a GC-MS, Varian 4000 to identify them. Both chromatographs were equipped with a HP-5 capillary column (30m x 0.25mm x 0.25μm) with injector in splitless mode, flame ionization detector, using helium as a carrier gas (24 cm.s^−1^). The column temperature was kept at 40°C for 1 min, increased to 150°C at a rate of 5°C per min and finally increased again at a rate of 20°C per min until reaching 250°C. Plant volatiles were identified by comparing their mass spectra and Kovat index (KI) using n-alkane (C7–C30) standards (Kovats, 1965) with those of the NIST08 library. Some compounds had their identity confirmed by comparison with available synthetic standards.

### Statistical analyzes

All data were submitted to analysis of variance followed by Tukey’s or t test in the software R (R Core Team 2017), considering the experimental design as completely randomized for all bioassays. Differences were considered statistically significant when the *p-value* <0.05. To quantify the bacterium by qPCR, the obtained data were log transformed to stabilize the variance. VOCs production data were log-transformed and Pareto-scaled before analysis using Metaboanalyst (Chong et al., 2019).

## RESULTS

### Development of the *Bacillus* sp. RZ2MS9 stable GFP-tagged strain

The efficiency of transformation of the PGPR *Bacillus* sp. RZ2MS9 using the pNKGFP plasmid was 8.0 × 10^3^ transformants.μg of plasmid DNA^−1^. The transformation was confirmed by PCR, which was performed with the specific primers for pNKGFP: PPNKF and PPNKRII. The electrophoresis gel showed the appropriate size of the amplicon (~360 pb) (Fig. 1). In addition, the same amplicon was not observed when we used RZ2MS9 wt DNA as template for PCR amplification.

**Fig. 1.**
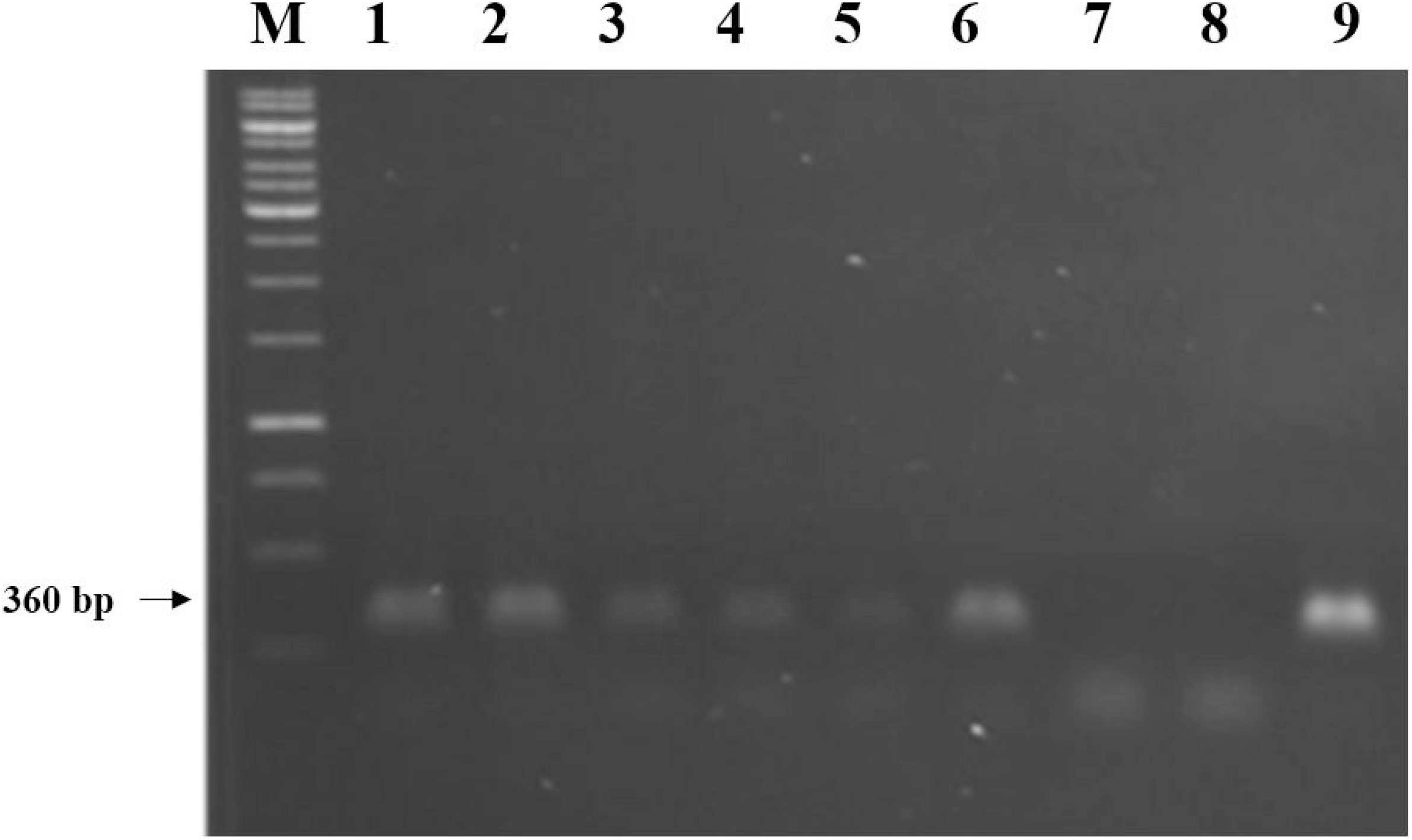
Molecular confirmation of the *Bacillus* sp. RZ2MS9 GFP-tagged transformants by PCR amplification using the primers PPNKF (5 ‘CCTTCATTACAGAAACGGC 3’) and PPNKRII (5 ‘GGTGATGCGTGATCTGATCC 3’). Lanes: **(M)** DNA ladder 1kb (Fermentas^®^); **(1-6)** RZ2MS9-GFP transformants; (**7)** RZ2MS9 wild type (negative control); **(8)** DNA-free water (blank control); **(9)** pNKGFP plasmid (positive control).

The measurements of bacterial growth revealed that GFP-tagged RZ2MS9 and RZ2MS9 wt presented the same growth curve pattern, even in the presence of the antibiotic (Fig. 2). This indicates that the insertion of the plasmid pNKGFP did not have any impact on the growth behavior of the bacterium. In all conditions, RZ2MS9 strains started the log phase approximately at 2.5 h h.a.i. and reached the stationary phase at 11 h.a.i..

**Fig. 2.**
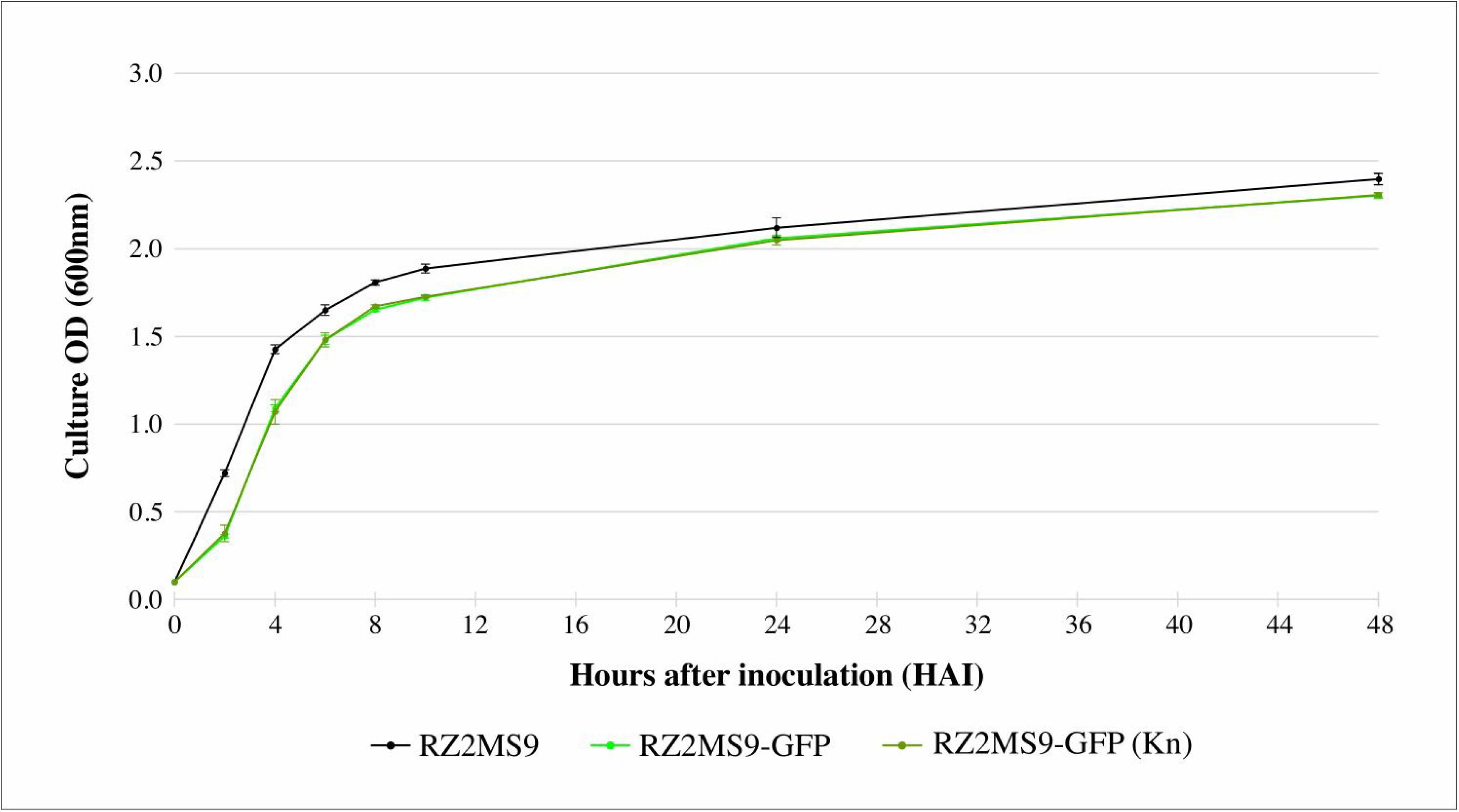
Bacterial growth curves of the RZ2MS9-GFP and of the RZ2MS9 wild type. All strains were grown in 50 ml of Luria Bertani (LB) broth medium, however the growth of the RZ2MS9-GFP was also evaluated in LB supplemented with kanamycin - kn (50 μg.ml^−1^). The bacterial growth, using four replicates per strain/condition, was estimated by measuring the Optical Density (OD) at 600nm every 2h, for 10h, and then at 24 and 48 hours after inoculation (h.a.i).

Finally, we found that the bacterial transformation had no influence on the ability of the *Bacillus* sp. RZ2MS9 to promote the growth of maize roots (Fig. 3). No statistically significant differences were detected between the treatment inoculated with RZ2MS9 wt and RZ2MS9-GFP for all of the evaluated parameters. On the other hand, both RZ2MS9 wt and RZ2MS9-GFP significantly improved the growth of maize roots when compared to the non-inoculated control for all root parameters evaluated (Fig. 3).

**Fig. 3:**
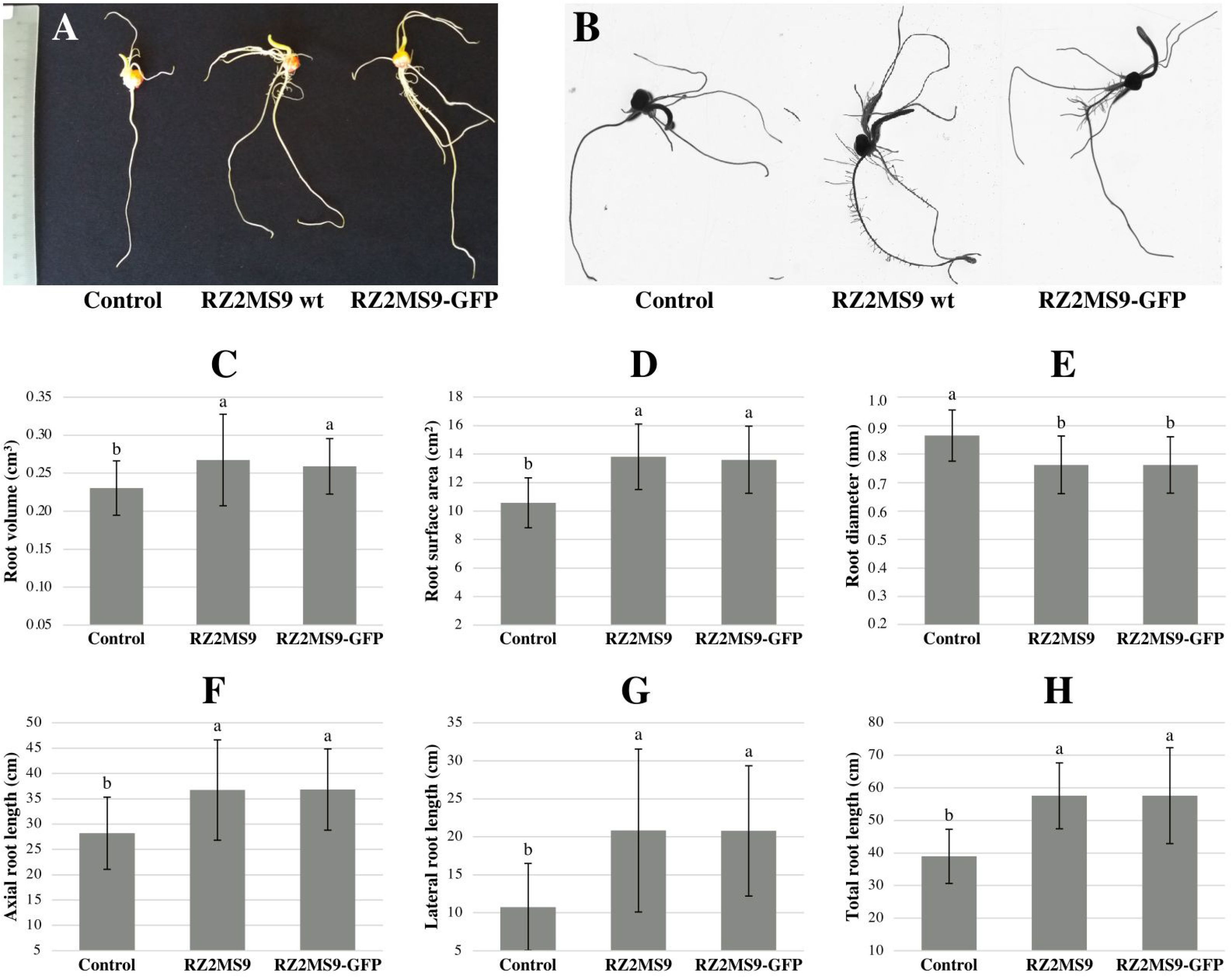
*In vitro* seed germination assay of maize (cultivar Dupont Pioneer^®^ P4285H) inoculated with RZ2MS9 wt and RZ2MS9-GFP. The germination assessment was performed 7 days after bacterial inoculation of maize seeds in wet germination paper towels **(A)**. The maize root system images were captured at 400 dpi resolution with the Epson^®^ Expression 11000XL scanner **(B).** The images were analyzed using the software WinRHIZO Arabidopsis, and the root parameters evaluated were: **(C)** root volume (cm^2^); **(D)** root surface area (cm^2^); **(E)** root diameter (mm); **(F)** axial root length (cm); **(G)** lateral root length (cm); **(H)** total root length (mm). Mean values (12 replicates per treatment) with the same letter are not significantly different (*P* > 0.05) according to Tukey’s test.

### Maize colonization by the GFP-tagged RZ2MS9

Using qPCR, we quantified the colonization of RZ2MS9-GFP in the maize’s roots and leaves, both at 15 and at 30 DAG. Overall, the number of bacterial cells detected at 30 DAG was lower than at 15 DAG, which could mean a decrease of maize colonization by RZ2MS9 over time (Fig. 4). The *Bacillus* sp. RZ2MS9-GFP was detected in leaf cells of inoculated plants, mostly in the chlorenchyma, in both palisade and spongy parenchyma. The tagged strain was also found colonizing the sub-stomata chamber, the epidermal cells and the xylem vessels of inoculated plants (Fig. 5B and D). A few RZ2MS9-GFP cells were found in maize root cells (Fig. 5F and H). On the other hand, no fluorescent GFP-tagged cells were observed in non-inoculated plants collected both at 15 and at 30 DAG (Fig. 5A, C, E and G). Therefore, the RZ2MS9-GFP was able to successfully colonize inner tissues of maize roots and leaves, demonstrating an endophytic behavior.

**Fig. 4.**
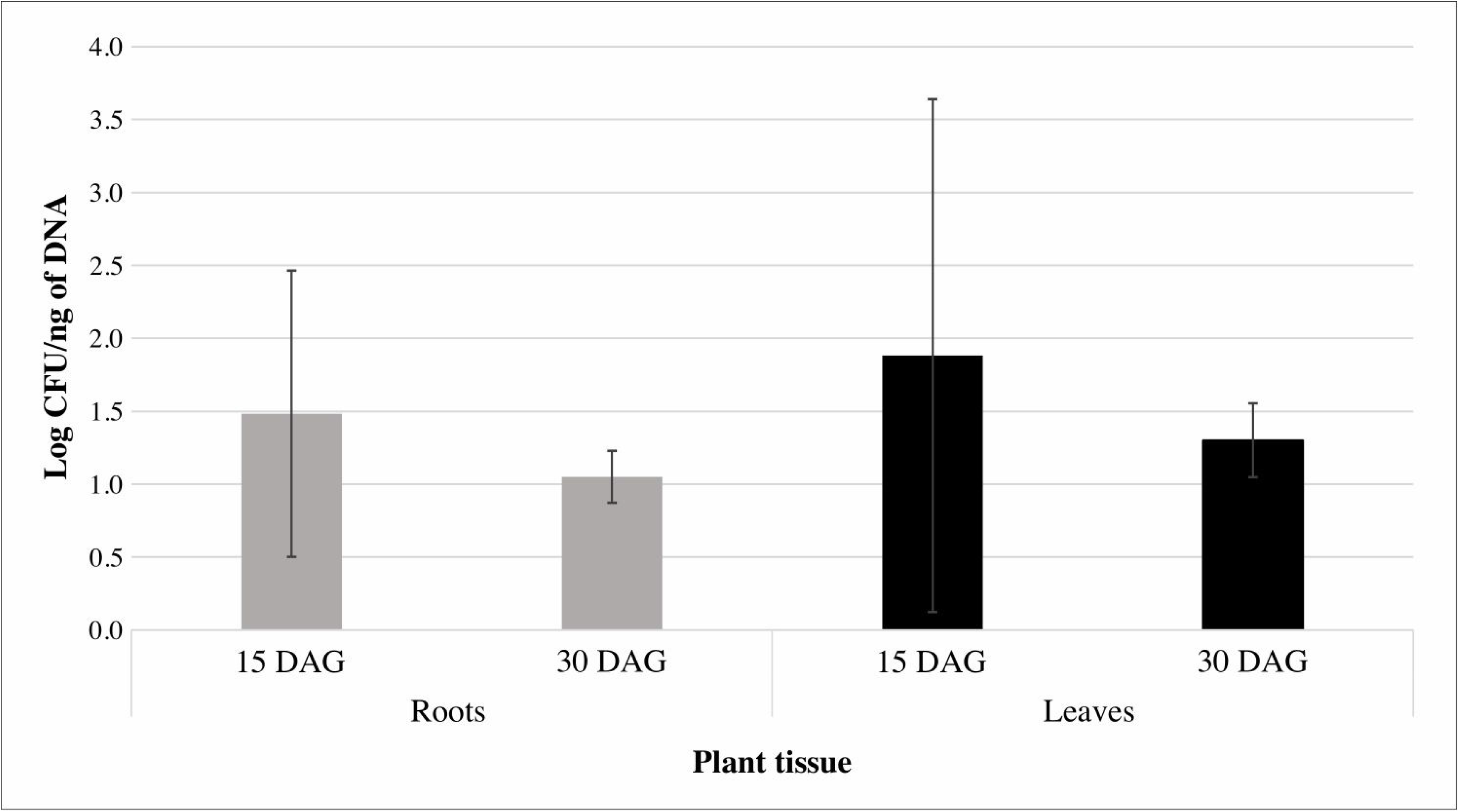
*Bacillus* sp. RZ2MS9-GFP abundance in different maize tissues during plant colonization. The bacterial cells were measured by qPCR at 15 and at 30 days after germination (DAG). The abundance data, in CFU/ng of DNA, were log-transformed to stabilize the variance. Data are presented as mean ± SE (*n*=3).

**Fig. 5.**
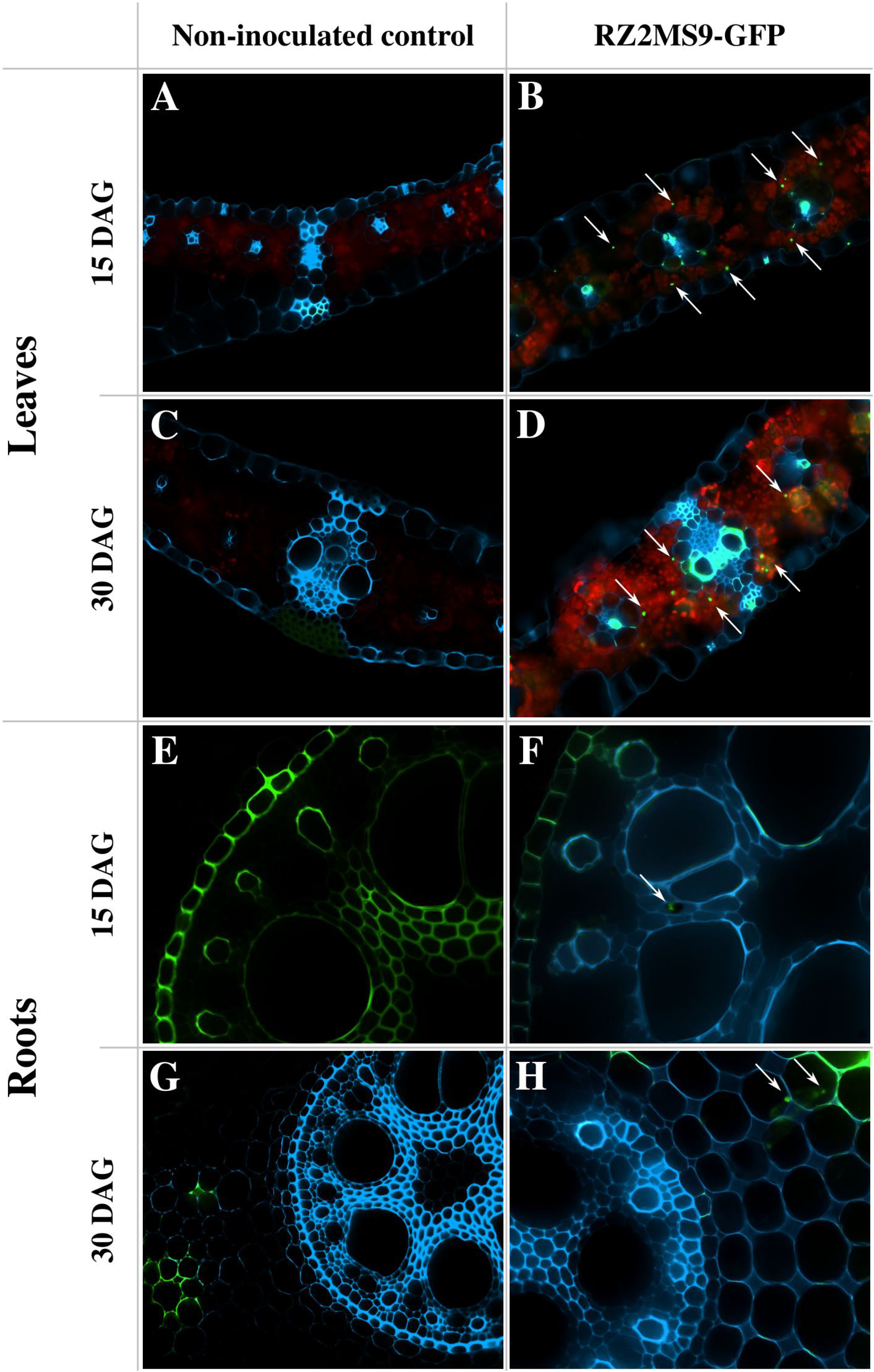
Fluorescence microscopy images of maize leaves and roots. The plants were inoculated or not with the *Bacillus* sp. RZ2MS9 tagged with green fluorescent protein (GFP)-expressing plasmid (RZ2MS-GFP). The transversal sections were performed in the tissues of maize collected at 15 and at 30 days after germination (DAG). The arrows indicate RZ2MS9-GFP cells in the leaves (**B and D**) and roots (**F and H**) of inoculated plants. No fluorescent bacterial cells were detected in the leaves (**A and C**) and roots (**E and G**) of non-inoculated control plants. The magnifications used were 100, 200 or 400 X.

### Maize growth promotion by the co-inoculation of RZ2MS9 and Ab-V5

Overall, the maize root system was positively affected by both treatments tested (RZ2MS9 alone and in co-inoculation with Ab-V5). However, the co-inoculation performed slightly better (Fig. 6). Root diameter and volume were only positively affected when co-inoculated with both strains. We observed increases of 12.5% and 33.9% in root diameter and volume, respectively, in plants treated with the combination of both strains (Fig 6D). The dry weight of maize roots was increased by 40.6% and 50.8% when inoculated with RZ2MS9 alone and when in co-inoculation with Ab-V5, respectively. For the dry weight of shoots, we detected an increase of 66.5% with the application of RZ2MS9 alone and of 79.6% with the co-inoculation. The RZ2MS9 alone and in combination with Ab-V5 increased maize shoots height by 22.3% and 20.2%, respectively. However, no significant differences among treatments were detected for root length, root projected area, root surface area, root length per volume, or stem diameter (Fig 6D).

**Fig. 6.**
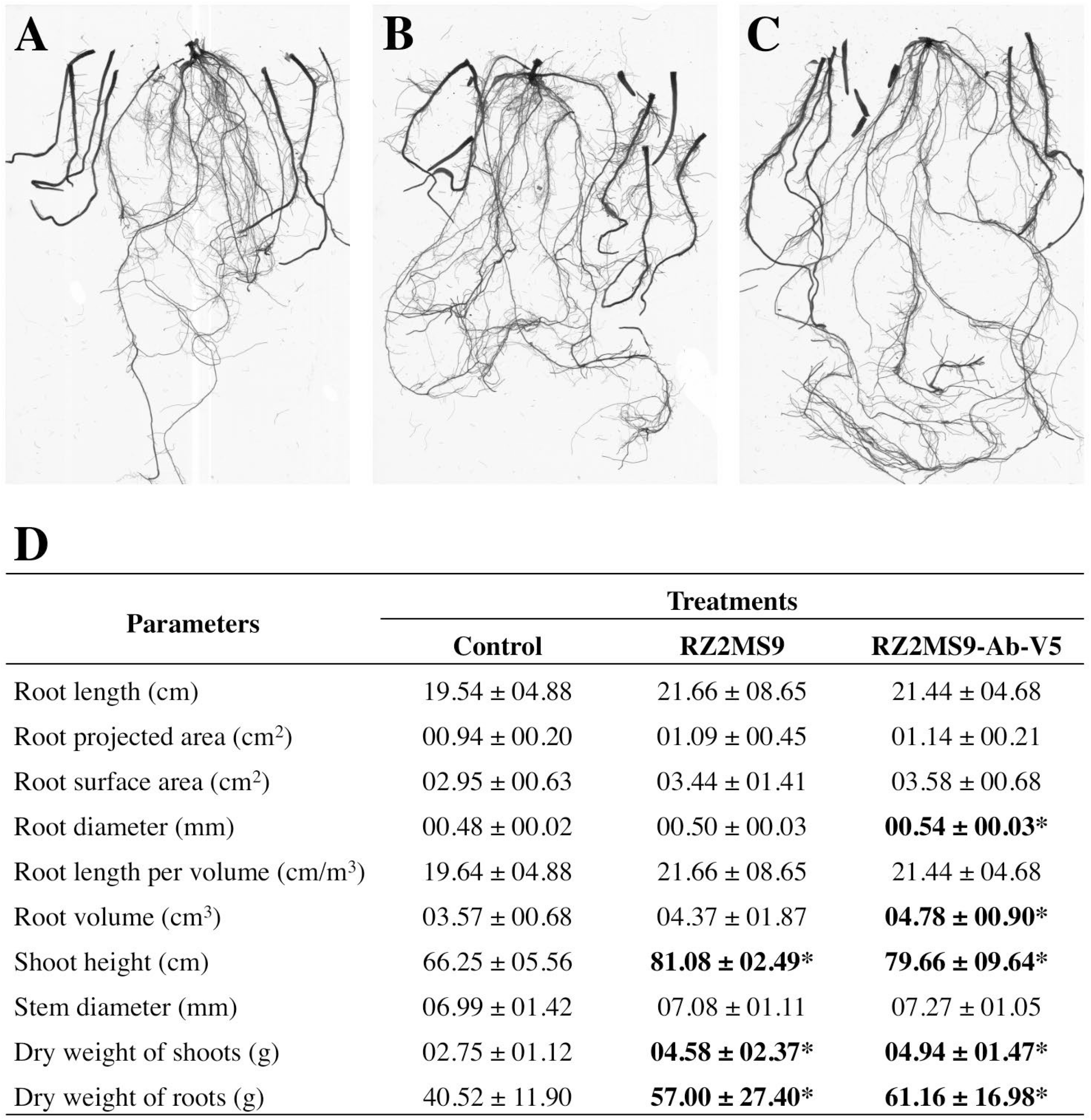
Effect on several growth parameters of 30-day old maize after inoculation with *Bacillus* sp. RZ2MS9 alone **(B)** and in co-inoculation with *Azospirillum brasilense* Ab-V5 **(C)**, in contrast with the non-inoculated control **(A)**. The data represent the means of 6 replicates per treatment ± the standard error. Asterisks indicate significant differences from the non-inoculated control according to the t-test (p-value <0.01) **(D)**.

### Inoculation effect on the production of plant Volatile Organic Compounds (VOCs)

Overall, inoculated maize plants produced more VOCs than non-inoculated plants. Maize plants inoculated only with RZ2MS9 and maize plants co-inoculated with RZ2MS9 and Ab-V5 produced 74.04 ng and 27.03 ng of metabolites.gram of dry plant tissue^−1^, respectively. Whereas non-inoculated maize plants produced 14.89 ng of metabolites per gram of dry plant tissue. A total of thirty-five VOCs were identified by GC, which were classified as aldehydes, alcohols, esters, hydrocarbons, monoterpenes, benzenoids and sesquiterpenes (Table 1). Twenty-seven metabolites significantly differed (*p-value* < 0.01) among the treatments. Eight of them were more abundant in maize plants that were co-inoculated with RZ2MS9 and Ab-V5: 2-hexanol, 3,4-dimethyl, dodecane, (E)-3-undecene, (E)-beta-farnesene, (E)-dodecen-1-ol, heptanal, indole and (Z)-3-hexen-1-ol. On the other hand, the inoculation with only RZ2MS9 enhanced alpha-cubebene production. When comparing inoculated plants (both co-inoculated and single inoculated) with the non-inoculated control plants, we found that the metabolites alpha-longipinene, alpha-ylangene, beta-linalol, decanal and nonane were more abundant in inoculated plants (Table 1).

**Table 1.**
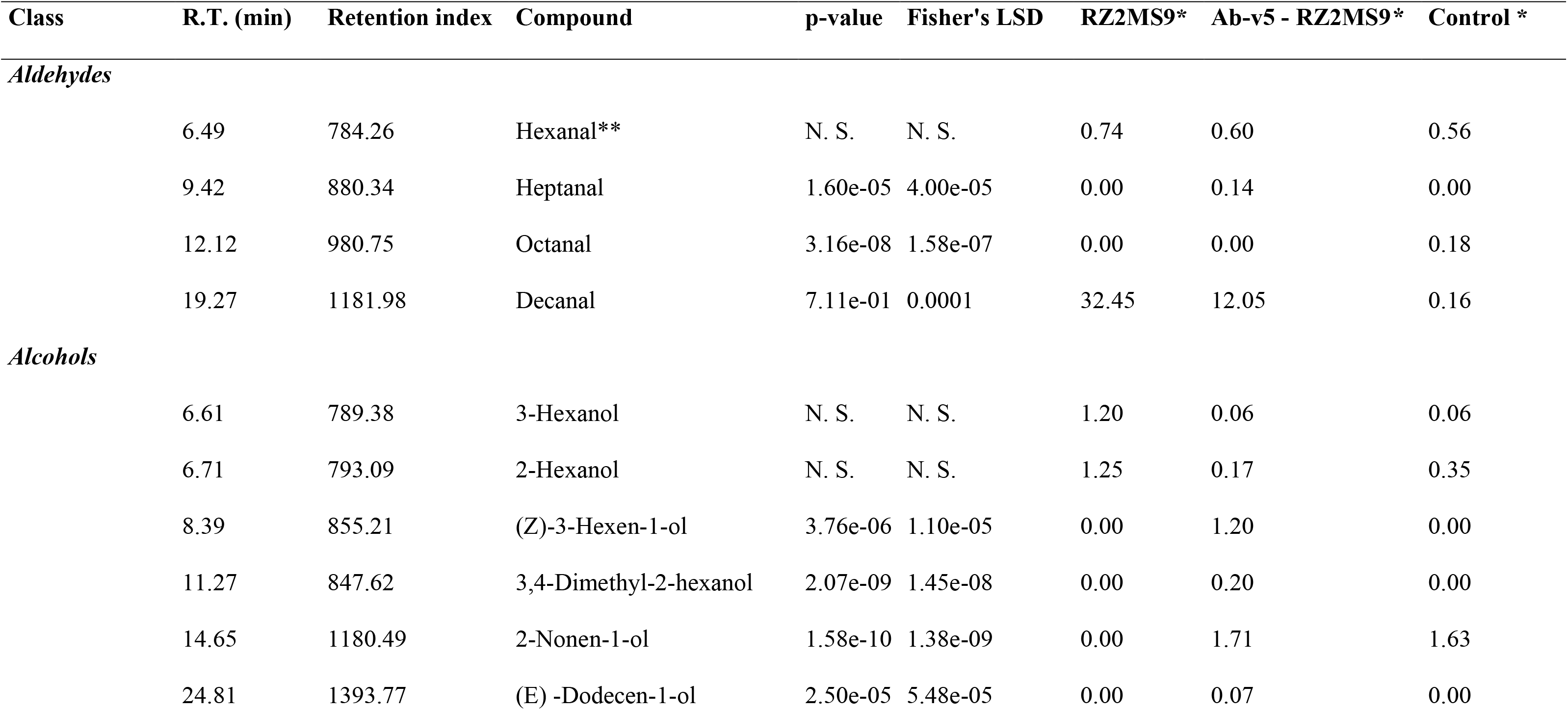
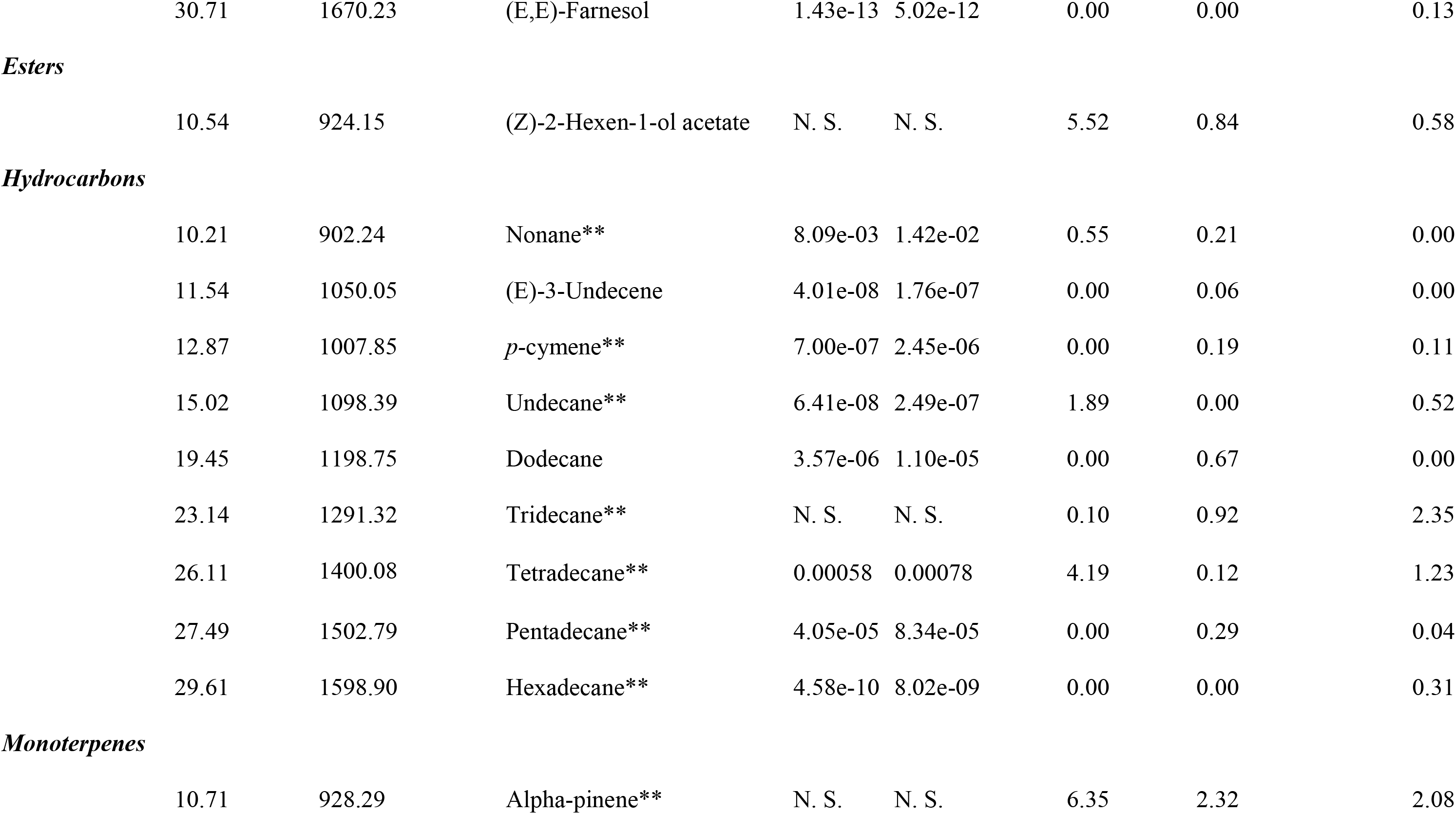
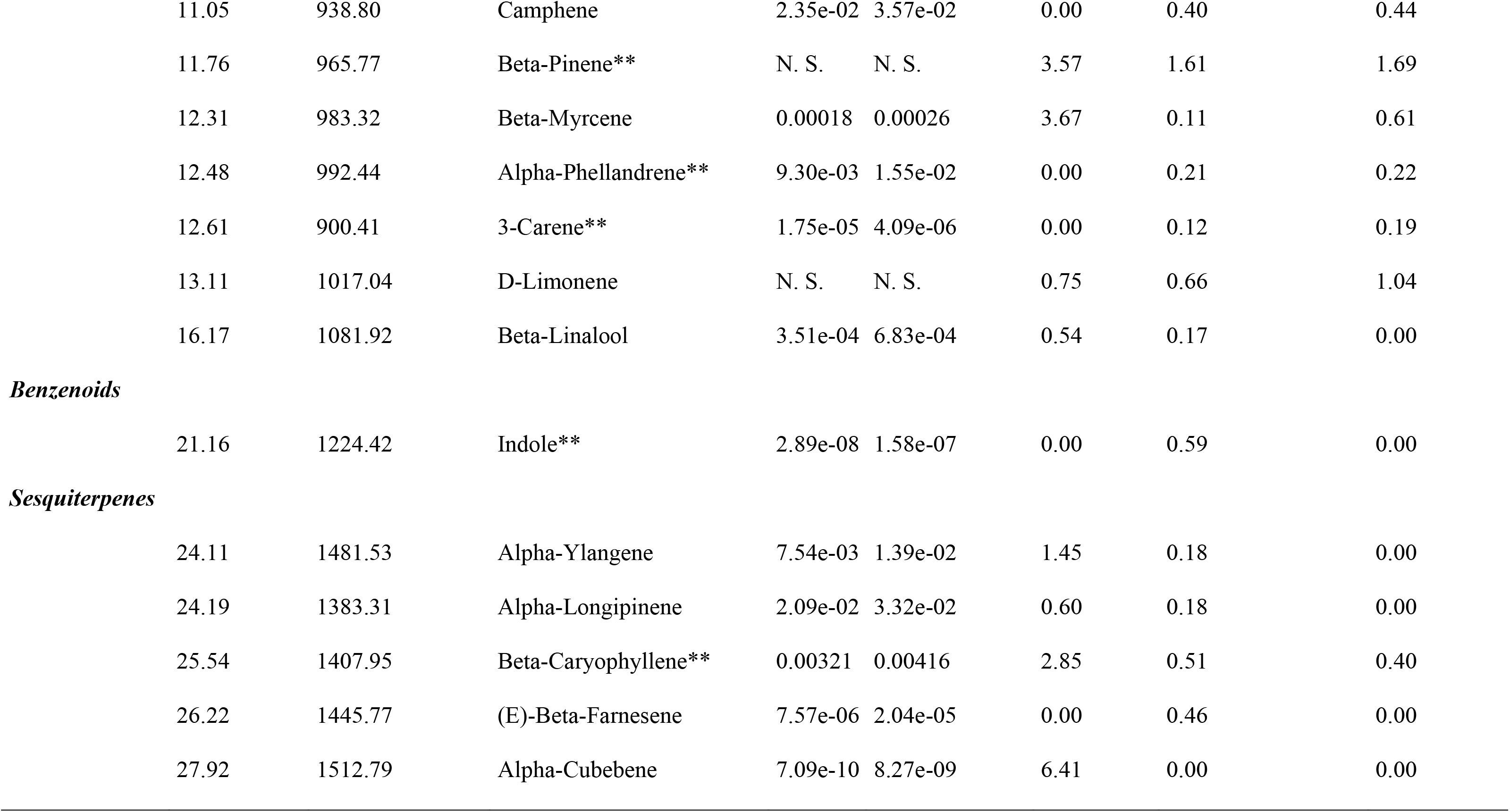
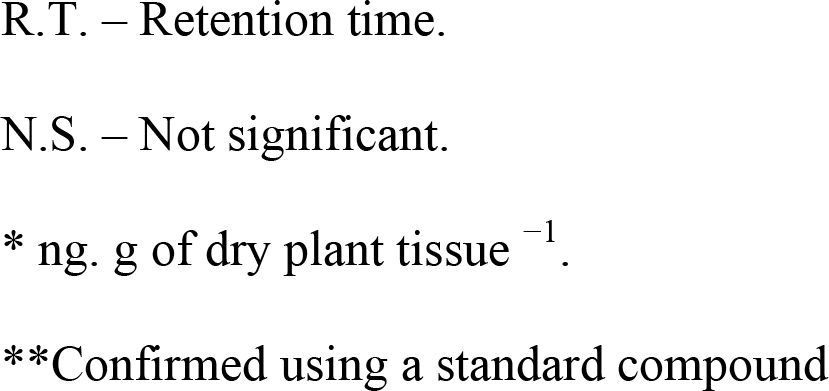
Volatile organic compound (VOC) production by maize single or co-inoculated with *Bacillus* sp. RZ2MS9 and *Azospirillum brasilense* b-V5

Interestingly, principal component analysis (PCA) significantly separated the VOCs profile emitted by co-inoculated maize plants from those emitted by RZ2MS9-inoculated plants and from non-inoculated control plants (Fig. 7).

**Fig. 7.**
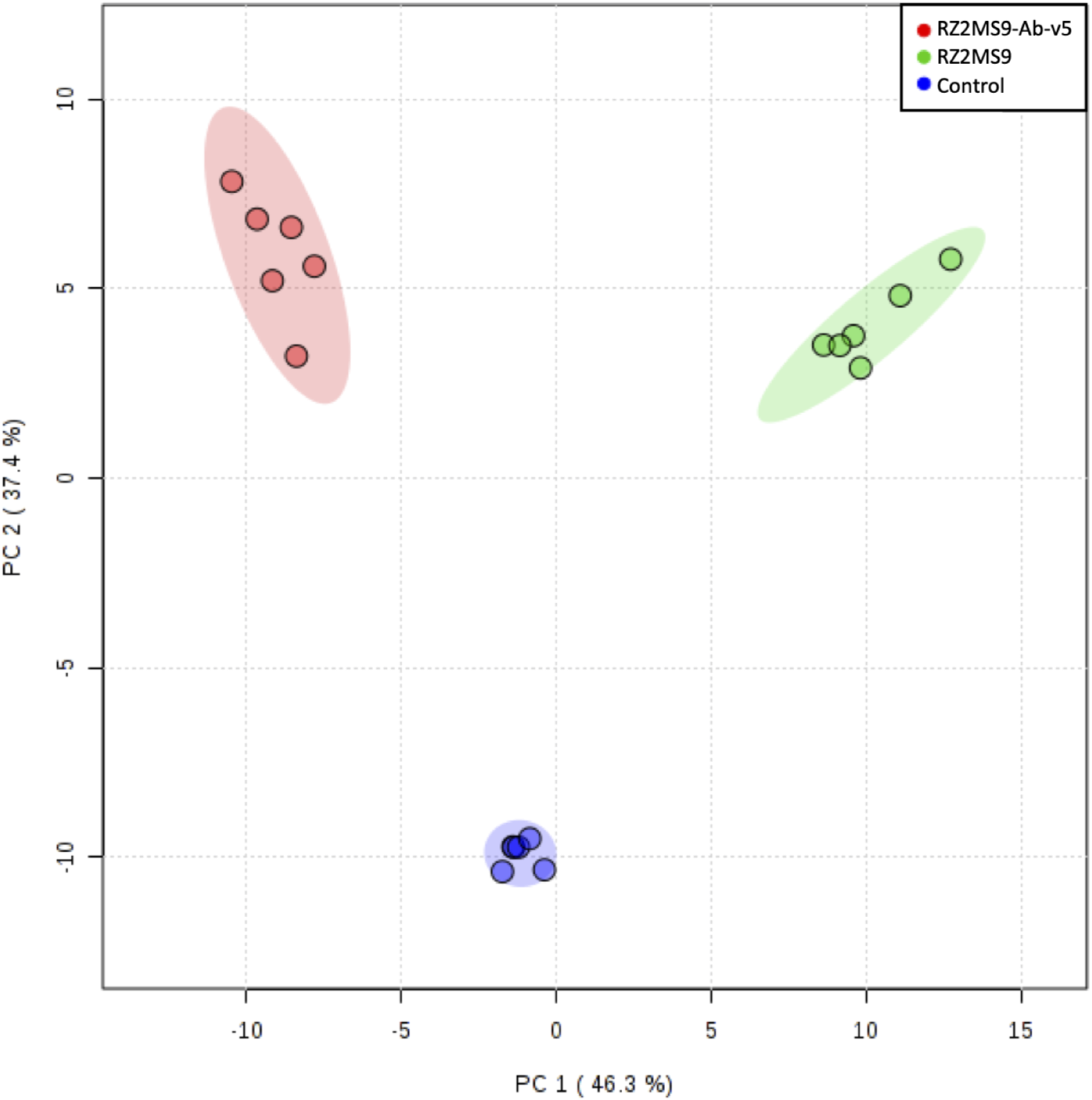
Principal component analysis of the volatile organic compounds (VOCs) emission patterns released by maize plants when inoculated with the combination of *Bacillus* sp. RZ2MS9 and *Azospirillum brasilense* Ab-V5 (RZ2MS9-Ab-V5) (in red); the *Bacillus* sp. RZ2MS9 alone (in green), and with no bacterial inoculation – control (in blue). Shaded areas represent the 95% confidence interval, and the explained variances are shown in brackets. 38

## DISCUSSION

The growing public concern on the use of chemicals in agriculture has increased the demand for efficient plant growth-promoting rhizobacteria (PGPR) as an alternative to synthetic fertilizers (Mendis et al., 2018). To be effective *in planta*, a PGPR candidate needs to be able to establish and maintain a sufficient population in the host plant (Krzyzanowska et al., 2012). Understanding the complex process of plant colonization by a PGPR is a big challenge and requires multiple approaches. However, it is a crucial step for the evaluation of a potential microbial inoculant.

In this study, we investigated the aspects of maize colonization by the *Bacillus* sp. RZ2MS9 using a polyphasic approach. Electrotransformation was the technique chosen to tag the RZ2MS9 with the *green fluorescent protein* (GFP). The transformation protocol used was efficient in inserting the integrative plasmid pNKGFP into the bacterial genome. This plasmid was first used to tag *Pantoea agglomerans* strain 33.1 in order to track its colonization in *Eucalyptus* seedlings (Ferreira et al., 2008). Later, this same tagged strain was monitored during the sugarcane interaction, showing a cross-colonization ability of the strain (Quecine et al., 2012). The bacterial pathogen *Leifsonia xyli* subsp. *xyli*, causal agent of ratoon stunting disease in sugarcane, was also transformed with the pNKGFP. The monitoring of the transformant Lxx::pNKGFP revealed some new colonization niches in sugarcane tissues by this pathogen (Quecine et al., 2016).

Here, we selected one transformant, named RZ2MS9-GFP, to be monitored during maize colonization. The tagged strain had only one integrative copy of the plasmid (data not shown), and measurement of bacterial growth showed that both GFP-tagged and the wild type (wt) RZ2MS9 exhibited the same growth curve pattern. It is known that the expression of new introduced genes may disturb normal cellular process (Wu et al., 2016). In fact, the endophyte *Pseudomonas putida* W619 presented a negative effect on poplar plants health and growth when GFP-labelled (Weyens et al., 2012). However, both the wild type and the GFP-tagged mutant of the diazotroph *Paenibacillus polymyxa* P2b-2R were able to promote the growth of pine (Tang et al., 2017), canola (Padda et al., 2016), and maize (Padda et al., 2017). In our work, a maize seed germination test was performed to ensure that the transformation did not affect the ability of RZ2MS9-GFP to promote the plant growth. We confirmed that both the RZ2MS9 wt and the RZ2MS9-GFP strains had the same performance when improving the maize root system growth as compared to the non-inoculated control.

The qPCR and fluorescence microscopy analyses revealed that the RZ2MS9-GFP was able to colonize internal tissues of maize plants, such as roots and leaves, demonstrating the bacterial endophytic behavior. Many efficient PGPR were reported inhabiting internal plant tissues (Hardoim et al., 2008). The *Burkholderia* sp. strain PsJN::*gfp2x* colonized *Vitis vinifera* L. cv. Chardonnay from the roots to the leaves (Compant et al., 2005). Since RZ2MS9 is a rhizobacteria (Batista et al., 2018), we hypothesis that the RZ2MS9-GFP penetrated the plant through the roots and migrated through the xylem to colonize the shoots and reach the leaves. Similarly, Hao and Chen (2017) GFP-tagged the PGPR *P. polymyxa* strain WLY78 and evaluated its colonization in maize. The authors observed that the strain was able to colonize the whole plant, detecting it in the cells of the roots, in the vascular system, and in the leaves.

Recent studies have shown that the adaptability and performance of bacterial inoculants could be improved by using mixed inoculants of multiple microbes, which are also known as microbial consortia (Sohaib et al., 2020). Here, we showed that the combination of RZ2MS9 with the *A. brasilense* strain Ab-V5 improved the effect of the *Bacillus* in promoting maize growth, suggesting a synergistic interaction between the tested strains. The co-inoculation was particularly effective in increasing root diameter and volume, which probably led to the overall increased dry weight of the maize shoots. Similarly, the co-inoculation of four PGPR, the *Pseudomonas putida* KT2440, the *Sphingomonas* sp. OF178, the *A. brasilense* Sp7 and the *Acinetobacter* sp. EMM02, demonstrated higher performance in promoting maize growth when compared to single and non-inoculated treatments (Molina-Romero et al., 2017). Cassán et al. (2009) observed that the co-inoculation of maize seeds with *A. brasilense* Az39 and *Bradyrhizobium japonicum* E109 resulted in the improvement of shoot length and of the shoot and root dry weight.

Different mechanisms can be involved in the plant growth-promotion triggered by a PGPR. In addition to all the previously demonstrated plant growth promoting traits displayed by the RZ2MS9, its positive effect on maize growth shown in the present study (both in the seed germination test and in the greenhouse assay) may be related with the carbon/nitrogen balance (Osuna et al., 2015). It is known that the carbon/nitrogen balance is crucial for the regulation of gene expression of pathways related to seed germination and plant development (Osuna et al., 2015). Several genes related to nitrogen metabolism were identified in the *Bacillus* sp. RZ2MS9 genome, among them the nitric oxide synthase oxygenase (*nos*), which catalyzes the production of nitric oxide (NO), and the nitrite transporter (*nirC*), which catalyzes the nitrite uptake and export across the cytoplasmic membrane (Bonatelli et al., 2020). The *A. brasilense* Ab-V5 also presents nitrogen fixation genes *nif* and *fix* which confer its ability to fix atmospheric nitrogen (Hungria et al., 2018). Moreover, both RZ2MS9 and Ab-V5 are indole-acetic acid (IAA) producers and carry genes related to the synthesis of this phytohormone in their genomes (Batista et al., 2018; Hungria et al., 2018).

In addition to the promotion of maize growth, the inoculation of RZ2MS9 alone and in combination with Ab-V5 altered the plant’s emission and composition of VOCs. Other works reported the same effect for different plant species when inoculated with PGPR (Santoro et al., 2015; Cappellari et al., 2017). The difference in plants’ VOCs emissions may have been caused by several reasons, such as the bacterial production of metabolites or the bacterial metabolization of VOCs produced by the plant, or it could be due to a change in the plant metabolism upon bacterial colonization (Ferré-Armengol et al., 2016).

The alteration in the plants’ VOCs emissions can even affect the way the plant interacts with its surroundings. Plants use VOCs to communicate with other plants, which is also a form of cross-kingdom communication (Farmer et al., 2001). Changes in secondary metabolites, such as in the VOCs, are also often involved in plant defense mechanisms. The inoculation of RZ2MS9 and Ab-V5, both in single and in co-inoculation, enhanced the production of different terpenoids, especially those belonging to the sesquiterpene and monoterpene class. Terpenoids are among the major constituents of plant’s VOCs emissions and they are related with indirect plant defense via tritrophic interactions (Das, 2013). The enhancement of emitted terpenoids upon PGPR inoculation is common (Banchio et al., 2009; Cappellari et al., 2013; Santoro et al., 2015).

We also observed that upon co-inoculation of RZ2MS9 and Ab-V5, the indole production was significantly enhanced when compared with the RZ2MS9 single inoculation and with the control. Indole is a benzenoid compound and a precursor of the amino acid tryptophan. Ballhorn et al. (2013) showed that rhizobia-colonized lima bean plants presented high production of indole and they hypothesized that the enhanced nitrogen availability from the rhizobia may be the reason for this. In fact, maize’s production of indole and other VOCs decreased upon the reduction of fertilization (Gouinguené and Turlings, 2002). In our work, the co-colonization may be playing an important role in maize plant nutrition and this will be further investigated.

Understanding the complex process of plant colonization by a PGPR is crucial to develop a more effective microbial inoculant for crops. This work not only increases knowledge about the interaction between *Bacillus* sp. RZ2MS9 and maize crops, but also provides a new possibility of its combined application with the commercial inoculant *Azospirillum brasilense*. The co-inoculation enhanced plant growth, favoring especially the root system, and altered the plants’ VOCs production. Future works will focus on understanding the molecular mechanisms involved in the interaction between the maize plant and this microbial consortium.

## ACKNOWLEDGEMENTS

This work was supported by grants from the São Paulo Research Foundation - FAPESP (Proc. No. 2015/01188-9 and 2014/50871-0) and National Council for Scientific and Technological Development - CNPq (Proc. No. 465511/2014-7). We would like to thank FAPESP for the fellowships granted to Jaqueline Raquel de Almeida (Proc. N. 2016/16868-8) and Rafaela Cristina dos Santos (Proc. No. 2016/1795-2). We also thank Coordination for the Improvement of Higher Education Personnel-CAPES for the fellowship granted to Maria Leticia Bonatelli (Proc. N. 88882.317565/2019-01) and CNPq for the fellowship granted to Carolina Alessandra de Almeida Hayashibara (Proc. N. 140587/2018-7).

